# Pupil size reflects the content of covertly attended afterimages

**DOI:** 10.64898/2026.05.04.722601

**Authors:** Ana Vilotijević, Sebastiaan Mathôt

## Abstract

Does attention operate within afterimages? Here we show that it does, using a novel pupillometry-based paradigm. Participants fixated centrally while bright and dark peripheral stimuli were presented, and a central cue directed attention to one of them. Over time, the stimuli perceptually faded due to adaptation and were then removed, leaving strong, negative afterimages. We found that pupil size tracked the brightness of the attended stimulus both during perceptual fading, when stimuli were present but perceptually weakened, and during perception of afterimages, when no physical stimuli were present. In the latter case, pupil size reflected the brightness of the negative afterimage rather than the preceding physical stimulus. This finding shows that covert attention can be directed within afterimages. More broadly, the results suggest that attention to afterimages bridges the gap between external and internal attention, challenging the notion of a strict dichotomy and supporting the view that this distinction is better understood as a continuum.

## Introduction

Afterimages are visual percepts that occur after viewing an adapting stimulus for a few seconds and then fixating on a neutral background (Daw, 1962; Anstis et al., 1978; Shimojo et al., 2001; Zaidi et al., 2012). For example, after staring at a white circle, you perceive an opposite or complementary percept (a black circle) once you look away. Afterimages are spatially localized, phenomenologically vivid, and in some cases strikingly stimulus-like, despite the absence of corresponding external input. But are afterimages also processed in the same way as external input, and thus susceptible to covert visual attention? Surprisingly, this fundamental question has never been conclusively answered, in part because it is methodologically challenging to do so. Here we use a recently developed pupillometric method (Vilotijević & Mathôt, 2024) to show that covert visual attention operates on afterimages.

Attention modulates sensory representations by enhancing or suppressing stimulus-driven activity, for example depending on task demands (Carrasco, 2011; Desimone & Duncan, 1995; Reynolds & Chelazzi, 2004). Very few studies have attempted to examine attention within afterimages (Bachmann & Murd, 2010), because it is difficult to assess directly. As a result, prior work mainly focused on manipulating attention to the adapting stimulus that precedes the afterimage (Brascamp et al., 2010; Lak, 2008; Lou, 2001; Suzuki & Grabowecky, 2003; Van Boxtel et al., 2010; Wede & Francis, 2007). These studies generally found that attending to the adapting stimulus reduces the strength or duration of the subsequent afterimage (but for the opposite results see Travis et al., 2017). However, such effects reflect how attention shapes the formation of afterimages, rather than how attention operates on the afterimage itself, which is the focus of the present study.

The most direct test of whether and how covert attention operates on afterimages comes from Bachmann and Murd (2010; in turn based on Lou, 2001). In their study, participants were presented with four colored disks for 24 s, after which the disks were removed, leaving four colored afterimages in their place. A color word (e.g. “red”) subsequently appeared, instructing participants to covertly search for the afterimage with the matching color. (There was no response associated with this visual-search task.) As soon as the first afterimage had faded, which typically happened after about four to eight seconds, participants indicated whether the first-to-fade had been the cued disk or any of the others. Crucially, participants indicated that the cued disk had faded first on about 50% of trials, far exceeding the 25% chance level. From this, Bachmann and Murd (2010) concluded that attention operates on afterimages, causing them to fade faster.

Bachmann and Murd’s (2010) conclusion was reasonable given the aforementioned well-established finding that covert attention to adapting stimuli also results in faster fading of subsequent afterimages. However, as they themselves mention, their results could also be explained by a response bias such that participants are more likely to over-report the cued stimulus due to other factors. As our results will show, this was likely the case.

Studying covert attention within afterimages is methodologically challenging because afterimages always reflect preceding adapting stimuli and cannot be directly controlled by a computer; simply put, it is not possible to present brief afterimage targets at cued or uncued locations within an afterimage display to test how fast and accurately these targets are detected. To address this limitation, we build on a pupillometry-based method that we recently developed (Vilotijević & Mathôt, 2024), which allows us to infer the allocation of covert attention within afterimages. In this paradigm, participants fixate centrally while two peripheral, fuzzy stimuli are presented, one bright and one dark. Participants covertly attend to one of the stimuli based on a central cue. Over time, the stimuli undergo perceptual fading as a result of adaptation. Once adaptation is complete, the stimuli are removed, leaving afterimages at the previously stimulated locations. We measure pupil size as an objective index of the perceived brightness of attended stimuli (Mathôt et al., 2013; Naber et al., 2013; Binda et al., 2013; Laeng et al., 2011; Mathôt, 2018; Vilotijević & Mathôt, 2023b). Because the pupil tracks not only physical luminance but also higher-level cognition—even in the absence of changes in external input—it provides a means to assess how attention modulates the perceptual content of afterimages. Accordingly, if afterimages are treated as external visual signals, attending to them should modulate pupil size in accordance with their perceived brightness. In other words, covertly attending to a dark stimulus should result in larger pupils than covertly attending to a bright stimulus. However, when the dark stimulus is removed, the effect should flip around: covertly attending to the resulting bright afterimage should result in *smaller* pupils.

## Methods

### Open Science Statement

All experimental methods and analyses were preregistered https://osf.io/29epv/. All experimental methods and analyses are available https://osf.io/x2q5n/.

### Apparatus and data acquisition

The experiment was programmed in OpenSesame (Mathôt et al., 2012) using PyGaze (Dalmaijer et al., 2014) for eye tracking. Stimuli were presented on a ROG Swift OLED PG27AQDM monitor, running at 240 Hz with a QHD resolution (Dimigen & Stein, in press). Pupil size and gaze position were recorded using an EyeLink 1000 eye tracker (SR Research Ltd., Mississauga, Ontario, Canada) with a sampling frequency of 1000 Hz.

### Participants

Data collection took place in the eye-tracking laboratory within the Department of Experimental Psychology at the University of Groningen. Our predetermined sample size consisted of 30 participants. Participants were students from the University of Groningen and received course credits for participation. Normal or corrected-to-normal vision was required for participation. On the basis of a checklist developed by the Ethical committee (EC-BSS) at the University of Groningen, the study was exempt from full ethical review (PSY-2324-S-0103).

### Procedure

Prior to the experiment, participants were seated approximately 60 cm from the display. A chin rest was used to stabilize head position, and an eye-tracking calibration and validation procedure were performed. In addition, a one-point drift correction was conducted before each trial. Trials were organized into blocks consisting of an adaptation period followed by five perceptual fading trials, and then a single afterimage trial. This sequence was repeated throughout the experiment (Fig. 1A).

**Figure 1.**
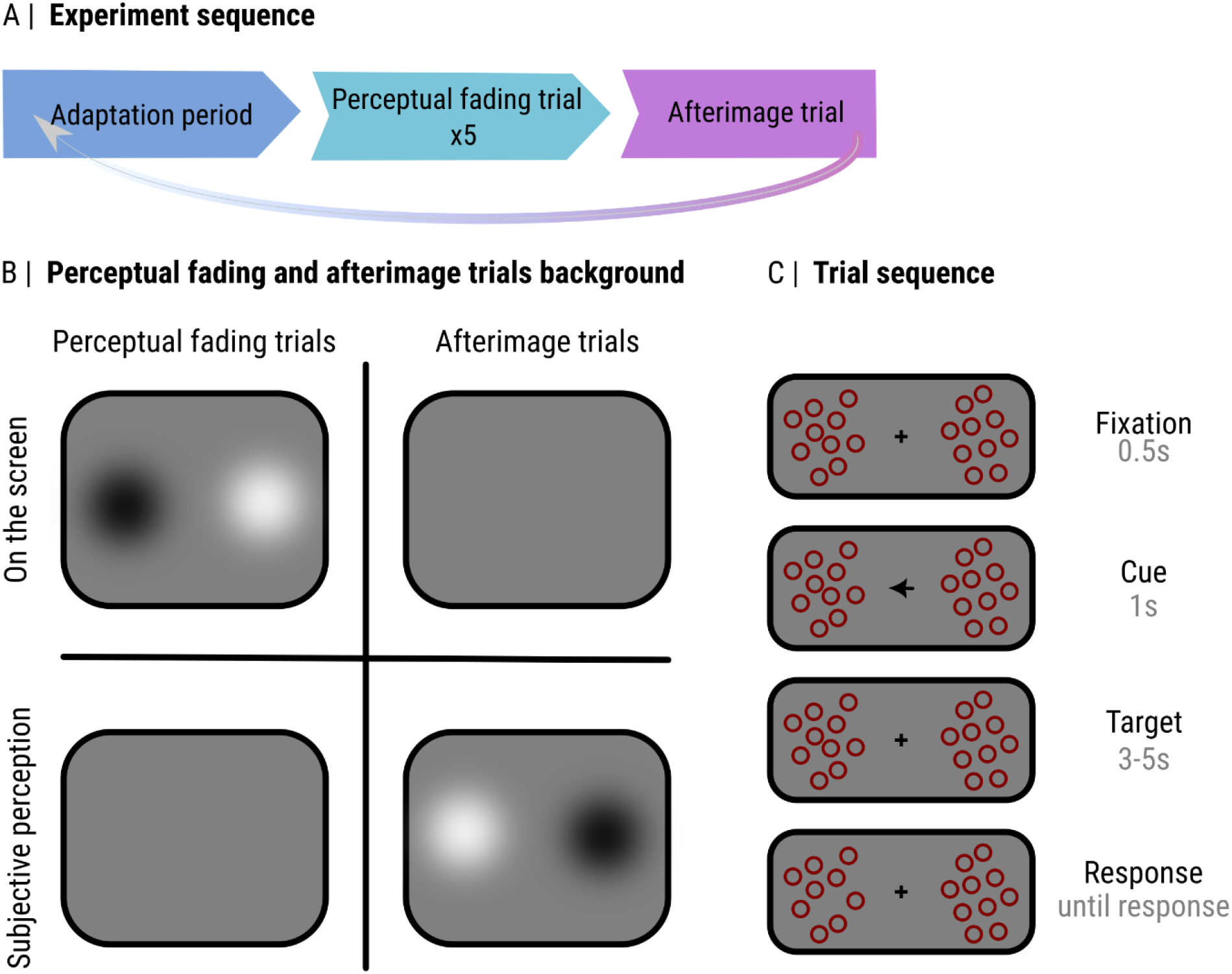
Experimental design and trial structure. **(A)** Experiment sequence. Each block began with the adaptation period, followed by five perceptual fading trials, and concluded with an afterimage trial. **(B)** Stimuli and subjective percepts in perceptual fading and afterimage trials. During perceptual fading trials, bright and dark patches were physically present on the screen but perceptually faded from awareness. During afterimage trials, the screen was uniform, and participants perceived negative afterimages at the previously presented stimuli. **(C)** Trial sequence. Each trial started with a fixation period (0.5 s), followed by a spatial cue (1 s) indicating the to-be-attended side. Next, a target display (3–5 s) was presented, during which pupil participants covertly attended to the cued side and one of the bubbles (a target) disappeared. Once participants detected the target they gave a keyboard response.

#### Adaptation period

The adaptation period served to establish the perceptual fading effect. Two circular patches were presented on a gray background (23.94 cd/m^2^): one white patch (SD = 5.77°; 99.60 cd/m^2^) and one black patch (SD = 5.77°; 0.14 cd/m^2^). The patches were positioned symmetrically on the left and right sides of the screen, with the center of each patch located 14.14° from the central fixation point. The adaptation period lasted 10s.

#### Perceptual fading trials

Perceptual fading trials were visually identical to the adaptation period (Fig. 1B), except that 10 randomly distributed unfilled circles (radius = 0.15°) with a maroon outline were superimposed on each patch. Each perceptual fading trial began with a central gray fixation dot presented for 0.5 s. This was followed by a central arrow cue pointing either left or right for 1 s. The cue indicated the side to which participants should covertly attend. After the cue, the display with the patches and circles remained on the screen. On 50% of trials, one randomly selected circle (the target) disappeared. The target offset occurred on the cued side on 80% of target-present trials and on the uncued side on the remaining 20%. The disappearance occurred at a random time between 3 and 5 s after stimulus onset. Participants were instructed to maintain fixation on the central dot throughout the experiment and to attend covertly to the cued side. Their task was to detect the disappearance of the target circle and report it as quickly as possible by pressing the space bar. On trials in which no target disappeared, participants were instructed to withhold a response (Fig. 1C).

#### Afterimage trials

Afterimage trials were identical to perceptual fading trials, except that the black and white patches were removed from the display, leaving a uniform gray screen (Fig. 1B). Because these trials followed a prolonged period of adaptation (the adaptation period and perceptual fading trials), participants experienced negative afterimages of the previously presented patches. These afterimage trials therefore allowed us to measure pupil responses while participants attended to perceptual content generated by afterimages rather than by physical luminance differences on the screen.

On both perceptual fading and afterimage trials, participants received feedback in the form of a green (correct) and red (incorrect) fixation dot after each trial. The experiment consisted of 20 practice trials followed by 400 experimental trials. To ensure that accuracy was maintained at approximately 75% across all conditions, we implemented a Single-Interval Adjustment-Matrix (SIAM) procedure (Kaernbach, 1990). We staircased the opacity of the circles based on the participant’s response: after a hit, the opacity of the offsetting circle was decreased for 1% (thus increasing task difficulty); after a miss, the opacity of the offsetting circle was increased for 3% (thus decreasing task difficulty); after a false alarm, the opacity of the offsetting circle was increased for 4% (thus increasing task difficulty); after a correct rejection, no adjustments were made (thus not changing task difficulty).

## Results

### Data preprocessing

#### Data exclusion

Practice trials were not analyzed. First, we checked whether participants made eye movements and excluded trials in which the deviation from the center of the screen was larger than 10° for more than 100 ms consecutively (615 trials excluded). We intentionally used a liberal exclusion criterion, because during afterimage trials (our primary focus) no stimuli were on the screen, and it was therefore not crucial that participants maintained steady fixation (anecdotally, fixation stability also seemed reduced on afterimage trials). Trials containing baseline pupil sizes of ±2 z-scores were excluded (865 trials). In total, 1489 trials (14.81%) were excluded from the data.

#### Pupil size

For the analysis of the pupil data, we followed the recommendations from Mathôt and Vilotijević (2022). We first segmented the pupil data into the time window of interest: from cue onset until four seconds later, which was the earliest possible moment of target presentation. Next, segments of data that contained eye blinks were reconstructed using the blinkreconstruct() function from Python DataMatrix. All pupil data was down-sampled from 1000 to 100 Hz. Finally, to reduce the effect of random fluctuations in pupil size, we applied baseline correction by subtracting the mean pupil size during the first 50 ms after the cue presentation from all pupil-size values (Mathôt & Vilotijević, 2022; Mathôt, 2018).

### Statistical analyses

#### General approach

For all analyses below, we ran a cross-validation in combination with linear mixed effects (LME) to localize and test effects (from the Python library time_series_test). This is a preferred approach in analyzing pupillary data that we explain in detail in Mathôt & Vilotijević (2022). We used four-fold cross-validation, such that 75% of the data was used as a training set and the remaining 25% as a test set. For all statistical analyses, we aimed to employ a maximal random-effects structure, including by-participant random intercepts and slopes for all fixed effects and interactions (Barr et al. 2013). However, due to convergence issues in some cases, we had to simplify the model structure in some cases. These adjustments are transparently reported below.

#### Covert attention modulates pupil responses to afterimages, and does so differently from perceptual fading

We tested whether covert attention to bright/dark modulates pupil size both in perceptual fading and afterimage trials. The model included Pupil Size (from the cue onset until the earliest target’s offset) as the dependent variable, Trial Type and Cued Brightness as fixed effects, and by-participant random intercepts and slopes. We found a main effect of Trial Type, indicating a stronger initial pupil constriction in the afterimage as compared to perceptual fading trials (*z* = 14.23, *p* < .001, tested at 100 ms), a main effect of Cued Brightness, indicating smaller pupils when covertly attending to dark stimuli (bright afterimage) for afterimage trials (*z* = −5.51, *p* < .001, tested at 800, 900 ms; this effect appears stable over time, see Fig. 2D), and an interaction, indicating that the effect of Cued Brightness differed between afterimage and perceptual fading trials (*z* = 3.46, *p* = 0.001, tested at 3200, 1000, 900, 1800 ms; Fig. 2A,B).

**Figure 2.**
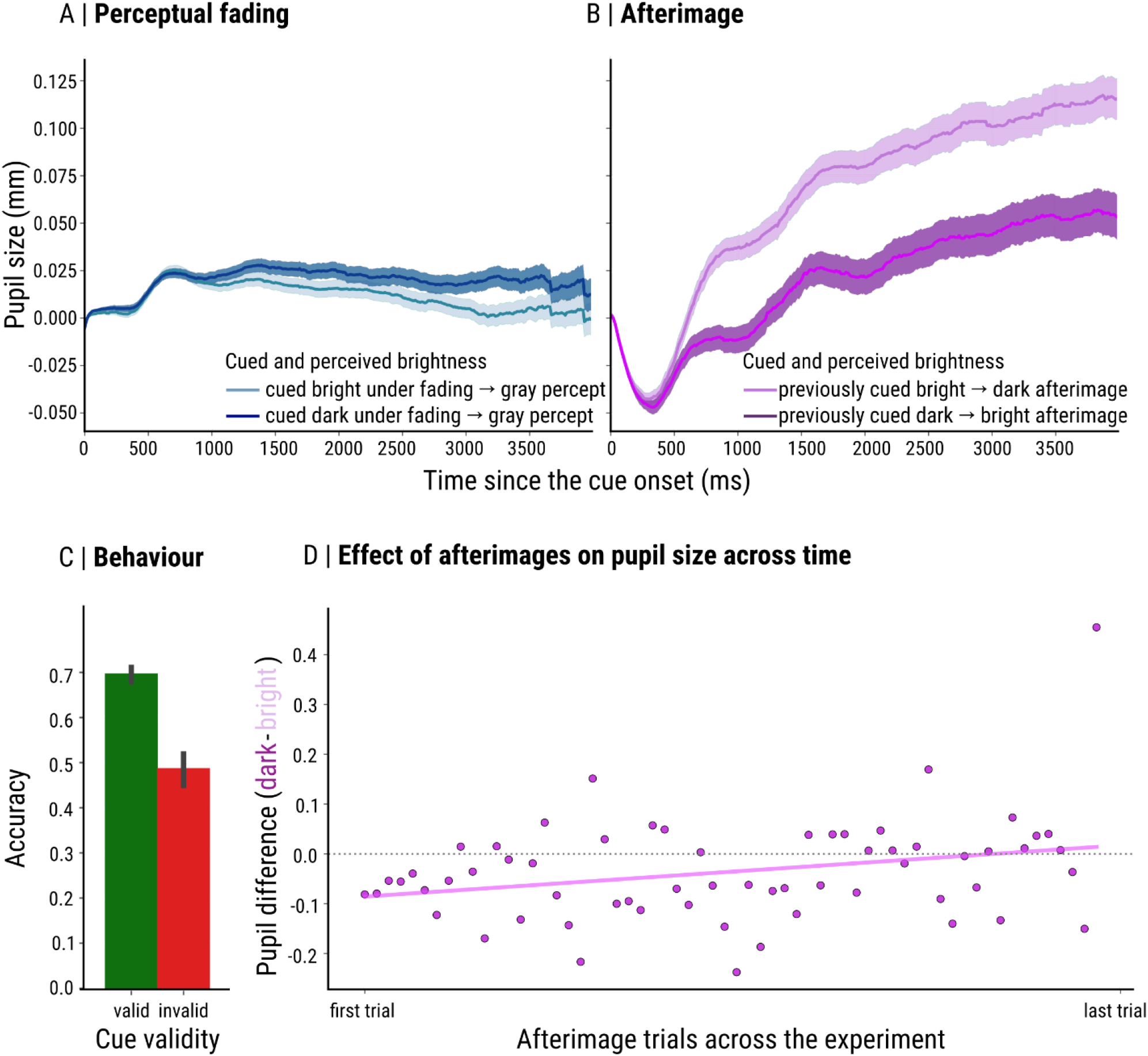
Covert attention modulates pupil responses during perceptual fading and afterimages. **(A)** Perceptual fading trials. Despite the fading of the stimuli from perceptual awareness, the pupil tracked the brightness of the attended stimulus, with larger pupils for dark as compared to bright stimuli. **(B)** Afterimage trials. Pupil size over time as a function of perceived afterimage brightness. Despite the absence of external input, the pupil tracked the brightness of the attended afterimage, with larger pupils when attending to dark as compared to bright afterimages. *Note*: Error bars represent the standard error. **(C)** Behavioral results. Accuracy as a function of cue validity. Performance was higher for valid than invalid trials, indicating that participants followed the cue and allocated attention to the instructed location. **(D)** Descriptive plot showing the pupil-size difference (dark − bright) across afterimage trials over the course of the experiment. The effect appears relatively stable over time.

#### Covert attention modulates pupil responses under perceptual fading

The analysis above used afterimages as reference Trial Type, and thus did not show directly whether covert attention to bright/dark modulates pupil size even when stimuli are faded away from perceptual awareness. To test this, we selected only perceptual fading trials, and tested the model that included Pupil Size (from the target onset^1^ until the earliest target’s offset) as the dependent variable, Cued Brightness as fixed effect, and by-participant random intercepts and slopes. We found a main effect of Cued Brightness, (*z* = 2.51, *p* = .011, tested at 2200, 2300 ms), indicating smaller pupils when covertly attending to bright stimuli under perceptual fading (Fig. 2A). This shows that pupil size is modulated by covertly attended brightness even when stimuli fade away from perceptual awareness, replicating our previous work (Vilotijevic & Mathôt, 2024).

#### Behavioral cueing effect

To assess the behavioral cueing effect, we fit generalized linear mixed models (GLMMs) with Accuracy as the dependent variable, Cue Validity as a fixed effect, and by-participant random intercepts. This analysis revealed a main effect of Cue Validity (*z* = 11.49, *p* <.001), indicating that participants attended the cued side of the display and were more successful in spotting the target offset from the validly cued side as compared to invalidly cued side (Fig. 2C). We next included Trial Type and its interaction with Cue Validity as additional fixed effects. This model again revealed a main effect of Cue Validity (*z* = 5.00, *p* < .001), but no main effect of Trial Type (*z* = −1.89, *p* = .057) and no interaction (*z* = 0.01, *p* = .990), indicating that the cueing effect did not differ between perceptual fading and afterimage trials.

## Discussion

Here, we show for the first time that covert attention can be directed within afterimages. We further show that this is reliably tracked by the pupil. These findings have two main implications: they extend our understanding of covert attention and clarify the mechanisms underlying cognitively driven pupil responses.

First, these results bear on theoretical accounts of what kind of representations are modulated by attention. According to influential frameworks, attention operates over both external sensory input and internally generated information, such as long-term memory (Chun et al., 2011; Kiyonaga & Egner, 2013; Myers et al., 2017; Nobre & Kastner, 2014; Van Ede & Nobre, 2023; Verschooren et al., 2019). However, this distinction may be less clear-cut than typically assumed (for a review see Oberauer, 2019). Afterimages occupy an intermediate position between external input and internally generated information: they are driven by prior external stimulation yet persist in the absence of ongoing input and are experienced as perceptual rather than mnemonic representations. The finding that attention can be directed within afterimages indicates that attentional selection extends to perceptual representations that are neither fully external nor fully internal. That is, attention to afterimages bridges the gap between external and internal attention, prompting us to reconsider the distinction between internal and external attention as a continuum rather than a dichotomy.

Second, our findings clarify the mechanisms underlying pupil responses. The pupil is known to respond not only to external sensory input, such as changes in luminance, but also to internally generated or maintained representations, such as those involved in attention, working memory, and mental imagery (Binda et al., 2013, 2014; Binda & Murray, 2015b; Bombeke et al., 2016; Mathôt et al., 2013; Unsworth & Robison, 2017; Blom et al., 2016; Vilotijević & Mathôt, 2023a,b; Fabius et al., 2017; Husta et al., 2019; Laeng & Sulutvedt, 2014; Naber & Nakayama, 2013; reviewed in Vilotijević & Mathôt, 2024). Extending this account, the present results show that pupil size also tracks afterimages, which—as mentioned—occupy an intermediate position. In other words, unlike luminance-driven responses, afterimages do not depend on current external input, and unlike mental imagery, they are not voluntarily generated, but arise spontaneously following sensory adaptation. This suggests that any form of perceptual experience—regardless of its source—engages common sensory mechanisms.

We have shown in our previous work (Vilotijević & Mathôt, 2024), and replicated here in the perceptual fading condition, that pupil size is modulated by covert attention to brightness even when stimuli fade from perceptual awareness. The magnitude of this effect is similar to that observed previously, with both effects on the order of ~0.02 mm. At first, this might seem to conflict with the finding that the pupil responds to perceptual experience in case of afterimages. However, we believe that the pupil does not track *only* physical input or *only* conscious perception, but rather the effective representation available to the visual system. One way to conceptualize this is in Bayesian terms, where internal and external signals are combined through a weighted integration process, with their relative contributions jointly determining a prediction of how much luminance is ‘out there’. That is, during perceptual fading, stimulus-driven signals continue to influence the pupil despite strongly reduced perceptual awareness, whereas in afterimages, perceptual representations influence the pupil despite the absence of input. In both cases, pupil responses track the representation that remains available—the visual system’s best guess of the outside world—regardless of whether the representation is primarily externally driven or internally sustained.

As mentioned in the introduction, Bachman and Murd (2010) reported that afterimages faded more quickly when covertly attended. How do their findings relate to ours? We believe the results of Bachman and Murd (2010) were due to a response bias: participants were more likely to indicate that a stimulus had faded from awareness when it had been cued. Using our method, we do not find any evidence that covert attention causes afterimages to fade more quickly. Instead, for the entire four second period that we measured, the brightness of attended afterimages strongly affected pupil size.

In sum, by showing that attention can be directed to afterimages and that this is reflected in pupil size, the present work demonstrates that perceptual representations—regardless of their origin—can guide both attentional selection and physiological responses. In this sense, the distinction between external input and internal percepts may be less clear than traditionally assumed. Instead, attention and pupil responses appear to operate over a shared representational space that includes externally driven and internally generated signals—and signals with properties of both.

## Transparency Statement

### Artificial Intelligence

No artificial-intelligence-assisted technologies were used in this research or the creation of this article.

### Ethics

The research protocol was approved for ethical considerations by the internal review board Ethical committee at the University of Groningen (PSY-2324-S-0103).

## Footnotes

1 This analysis deviates from the preregistration: the cue period (1 s) was excluded, as the effect of interest developed only after this interval and including it did not provide a meaningful test of the hypothesis.

